# Med12 and Med13 prevent tumorigenic dedifferentiation of intermediate neural progenitors and premature loss of neural stem cells

**DOI:** 10.64898/2026.03.11.711102

**Authors:** Rui Chen, Xiaosu Li, Wenwen Lu, Yanjun Hou, Sijun Zhu

## Abstract

Med12 and Med13 are components of the kinase module of the mediator complex. Mutations of Med12 and Med13 have been associated with neurodevelopmental disorders and various cancers. However, their functions in neural development are not well understood. Here we show that in the developing *Drosophila* brain, Med12 and Med13 are required to prevent tumorigenic dedifferentiation of intermediate neural progenitors (INPs) and maintain neural stem cell (NSC) self-renewal. We further demonstrate that Med12 and Med13 prevent INP dedifferentiation by coordinating with a subset of core mediator complex subunits to mediate the activation of genes required for INP fate commitment. In contrast, during the maintenance of NSC self-renewal, Med12 and Med13 antagonize the function of a different subset of core mediator complex subunits. Together, our findings reveal that Med12 and Med13 perform two distinct functions in neural progenitors by coordinating with one subset of core mediator complex subunits while antagonizing another.

**Highlights:**

- Loss of Med12 and Med13 causes dedifferentiation of intermediate neural progenitors
- Med12 and Med13 mediate the activation of target genes of PntP1
- Loss of Med12 and Med13 leads to premature loss of neural stem cells
- Med12 and Med13 act with one subset of core mediator subunits but oppose another

**eTOC blurb:** Zhu and his colleagues show that Med12 and Med13 promote cell fate commitment of intermediate neural progenitor cells and self-renewal of neural stem cells. Med12 and Med13 perform these two distinct functions by coordinating with one subset of core mediator complex subunits while opposing another to regulate the expression of different target genes.

## Introduction

Med12 and Med13 are two subunits of the mediator complex that regulates the expression of target genes of sequence-specific transcription factors by bridging the transcription factors with the general transcriptional machinery. The mediator complex is composed of the kinase module and the core mediator complex, the latter of which is composed of over 20 subunits and can be further divided into head, middle, and tail modules. Med12 and Med13 together with CDK8 and CycC comprise the kinase module that regulates the activity of the core Mediator complex ^1,2^. However, MED12 and MED13 can also act independently of the kinase module or the core Mediator complex ^3–6^. Unlike the core Mediator complex, which is nearly ubiquitously required for regulating gene transcription, MED12 and MED13 appear to mediate the expression of target genes of specific transcription factors as indicated by a relatively small number of genes regulated by MED12 and MED13 in a yeast microarray analysis ^7^. MED12 and MED13 regulate the development of various tissues and organs, such as the eye and wing disc in *Drosophila* and brain and neural crest in zebrafish, by modulating the expression of target genes of different signaling pathways ^5,6,8–10^. Importantly, in humans, mutations in MED12 or MED13 have been implicated in various developmental disorders, such as FG and Lujan syndromes, which share common clinical phenotypes including intellectual disability and macrocephaly, indicating their important roles in human neural development ^11–13^. In addition, high frequency of mutations in MED12 has been observed in a variety of tumors ^14–17^ and reduced adenosine-to-inosine RNA editing of MED13 in human brain cancers has also been reported ^18^, suggesting potential involvement of MED12/MED13 in tumorigenesis. However, the exact roles of Med12 and Med13 in neural development, especially in neural stem cells, are not well known.

Drosophila neuroblasts (NBs, the Drosophila neural stem cells) in the developing Drosophila central brain have provided an excellent model system for studying neural stem cells. There are two different types of NBs, type I and type II (Figure 1A, left). There are about 90 type I NBs and 8 type II NBs in each brain lobe ^19–22^. Type II NBs can be distinguished from type I NBs by their lack of the expression of the proneural protein Asense (Ase) ^20,23,24^. A single type II NB can generate several-fold more neurons with diverse types than a single type I NB because, unlike type I NBs that produce ganglion mother cells (GMC), type II NBs generate intermediate neural progenitor cells (INPs) ^19–21^. A GMC only divides once to produce two neurons, whereas an INP can undergo several rounds of self-renewing divisions like type I NBs to produce GMCs ^19–22^. Newly generated INPs are immature (thus called immature INPs or imINPs) and undergo a differentiation process to become mature INPs (mINPs) before they start to divide ^20,25^. imINPs are prone to dedifferentiate back into type II NBs before they are fully committed to the mature INP fate (Figure 1A, right) ^20,26,27^. Both types of NBs in the central brain continuously undergo asymmetric self-renewing divisions to produce neurons from late embryonic stages to early pupal stages except a period of quiescence from the end of embryonic stages to early larval stages. At early pupal stages, all NBs except the mushroom body NBs terminate their self-renewal ^22,28–31^. This termination is largely regulated by the mediator complex and temporal identity factors such as Caster, Seven-up, RNA binding proteins IGF-II mRNA-binding protein (Imp) and Syncrip/hnRNPQ (Syp), which inhibit cell growth and promote Prospero (Pros)-mediated differentiation and cell cycle exit of NBs. ^30,32,33^.

**Figure 1.**
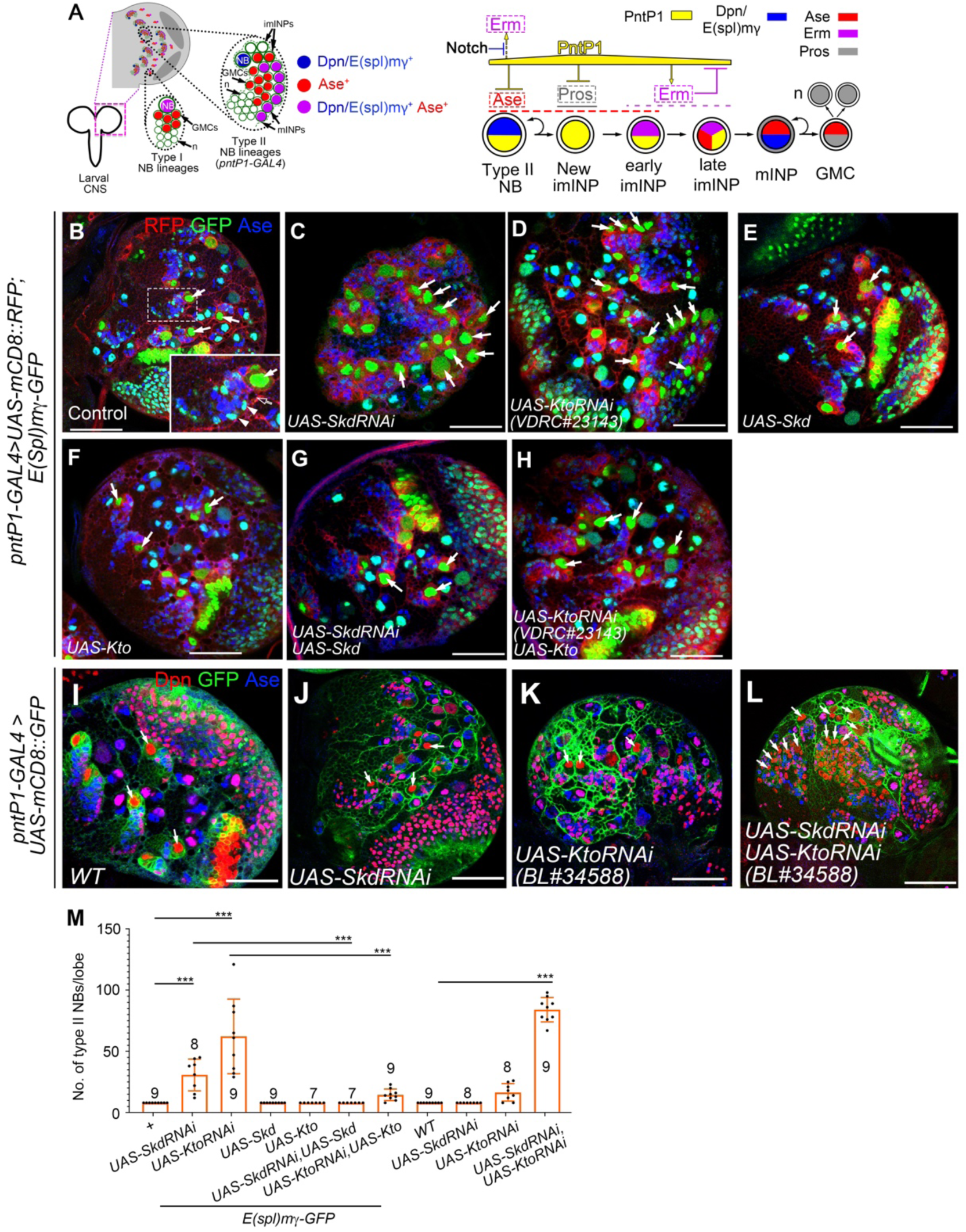
Skd or Kto is required to prevent the generation of supernumerary type II NBs. All images are from 3^rd^ instar larval brains with the *E(spl)mγ-GFP* transgene. Type II NB lineages are labeled with mCD8-RFP (B-H) or mCD8-GFP (I-L) driven by *pntP1-GAL4*. Brains are counterstained with anti-Ase and anti-Dpn antibodies. Examples of type II NBs are pointed with solid white arrows. Scale bars equal 50µm. (A) (Left) A diagram of a 3^rd^ instar larval brain lobe with both type I and type II NB lineages. (Right) A schematic diagram of the development of the type II NB lineage and some of the key genetic programs involved in type II NB lineage development. NB: neuroblast; imINP: immature INP; mINP: mature INP; GMC: ganglion mother cell; n: neuron. (B) A wild type brain lobe with the *E(spl)mγ-GFP* transgene contains eight type II NB lineages (only seven are shown). An enlarged view of the lineage highlighted with a dashed rectangle is shown in the inset. Arrowheads point to mINPs and the open arrow to imINP. (C-D) Knockdown of Skd (C) or Kto (D) in type II NB lineages in the presence of *E(spl)mγ-GFP* results in supernumerary type II NBs. (E-F) Expressing *UAS-Skd* (E) or *UAS-Kto* (F) in type II NB lineages does not affect the number of type II NBs. (G-H) Expressing *UAS-Skd or UAS-Kto* largely suppresses the generation of supernumerary type II NBs caused by knockdown of Skd (G) or Kto (H), respectively. (I) A wild type brain lobe without the *E(spl)mγ-GFP* transgene contains 8 type II NBs. (I-L) Knockdown of Skd (J) or Kto (K) alone in the absence of *E(spl)mγ-GFP* results in no or a very weak supernumerary type II NB phenotypes, respectively, but simultaneous knockdown of Skd and Kto induces a strong supernumerary type II NB phenotype (L). (H) Quantifications of number of type II NBs in brains with indicated genotypes. Data represent Mean ± SD. ***, *P* < 0.001. The number on each bar indicates the number of brain lobes examined.

The specification of type II NBs and the development of INPs are primarily regulated by the Ets family transcription factor Pointed P1 (PntP1) (Figure 1A, right) ^34^. PntP1 is specifically expressed in type II NBs and imINPs but not in type I NB lineages. In type II NBs, PntP1 specifies the NB identity by activating Tailless (TLL), which in turn suppresses Ase expression ^34–37^. In newly generated imINPs, PntP1 together with Buttonhead (Btd) suppress the expression of homeodomain protein Pros to prevent imINPs from prematurely differentiating into GMCs ^38,39^. Later during imINP development, PntP1 activates the expression of the zinc-finger protein Earmuff (Erm), which prevents dedifferentiation of imINPs by promoting INP maturation and cell fate commitment ^26,40^. Erm does so by acting together with Six4 to provide a feedback inhibition on PntP1’s activity and expression ^25,41,42^. Although PntP1 is both necessary and sufficient for the specification of type II NBs and the generation of INPs, maintaining the identity and self-renewal of type II NBs also requires Notch signaling. Notch activates the expression of bHLH family proteins such as E(Spl)mψ and Dpn, which in turn prevent precocious activation of Erm by PntP1 in type II NBs ^25,28,41,43–45^. In imINPs, termination of Notch signaling by Numb and degradation of Notch targets E(Spl)mψ, Dpn and other self-renewing factors allow Erm to be activated by PntP1 to promote INP maturation ^20,25,46–48^. In addition to these genetic programs, various epigenetic programs mediated by the SWI-SNF complex, Trithorax, histone deacetylase 1 (Hdac1/Rpd3), Hdac3, Brahma, etc. also play crucial roles in the development of INPs by regulating the chromatin status and coordinating with the genetic programs ^25,27,37,39,49^.

In this study, we investigated the role of Skd and Kto in *Drosophila* larval NBs. We found that Skd and Kto have two distinct roles. One is to regulate the specification of type II NBs and the cell fate commitment of INPs by mediating the activation of target genes of PntP1. Another is to maintain the self-renewal of larval NBs. Interestingly, Skd and Kto perform these two distinct functions by interacting with different subsets of subunits of the core mediator complex. In the cell fate commitment of INPs, Skd and Kto likely coordinate with one subset of core mediator complex subunits. Conversely, in maintaining NB self-renewal, Skd and Kto likely antagonize the function of a different subset of core mediator complex subunits that normally promote cell shrinkage and differentiation during early pupal stages. Together, our work reveals novel functions of Skd and Kto in neural stem cells, which may help us understand pathogenesis of various neural developmental disorders resulting from mutations in human Med12 and Med13.

## RESULTS

### Skd and Kto act synergistically to prevent the generation of supernumerary type II NBs

To identify novle transcription factors that regulate type II NB lineage development, we conducted an RNAi knockdown screen of transcription factors in type II NB lineages. In this screen, individual type II NB lineages are labelled with *UAS-mCD8-RFP* driven by type II NB lineage-specific *pntP1-GAL4* ^34^, which is also used to drive the expression of *UAS-RNAi* transgenes in type II NB lineages. Meanwhile, all NBs and mature INPs were labeled by E(spl)mγ-GFP ^46^. With these markers, we could identify type II NBs (large E(spl)mγ-GFP positive cells), imINPs (next to the type II NB but E(spl)mγ-GFP negative), and mINPs (small E(spl)mγ-GFP positive cells) in individual type II NB lineages (Figure 1B). Including a copy of *E(spl)mγ-GFP* transgene also makes animals more susceptible to the generation of supernumerary type II NBs ^42^, thus providing a sensitized background for identifying genes that suppress the generation of supernumerary type II NBs. In wild type larvae, each brain lobe contains 8 type II NB lineages with each lineage comprising one NB, 3-4 imINPs and 20-30 mINPs ^19–21,34^. We found that knockdown of Skd or Kto resulted in an average of 30 or 60 type II NBs per brain lobe, respectively (Figure 1C-D, M). The supernumerary type II NB phenotype resulting from Skd or Kto knockdown could be largely rescued by expressing *UAS-Skd* or *UAS-Kto* (Figure 1E-H, M), indicating that the supernumerary type II NB phenotypes resulting from Skd or Kto RNAi were not caused by off-target effects. These results suggest that Skd and Kto normally prevent the generation of supernumerary type II NBs.

Since Skd and Kto function together in the same kinase module of the mediator complex to regulate various developmental processes ^5,8,50^, we next investigated whether they also function together in type II NB lineages by testing if simultaneous knockdown of Skd and Kto synergistically enhances the supernumerary type II NB phenotype. Since inclusion of the *E(spl)mγ-GFP* could enhance the supernumerary type II NB phenotype as we showed previously, we knocked down Skd and/or Kto in absence of E(spl)m*γ*-GFP for testing their genetic interactions. Our results showed that in the absence of *E(spl)mγ-GFP*, knockdown of Skd alone did not cause production of any extra type II NBs, whereas knockdown of Kto alone only slightly increased the total number of type II NBs to 16 per lobe on average (Figure 1I-K, M). However, double knockdown of Skd and Kto synergistically increased the number of type II NBs to more than 80 per lobe (Figure 1L-M). The synergistic effect of Skd and Kto double knockdown support that they function together to prevent the generation of supernumerary type II NBs.

### Kto and Skd are ubiquitously expressed in the *Drosophila* central brain

Next we examined if Skd and Kto are expressed specifically in type II NBs or ubiquitously expressed in fly larval brains. For examining Kto expression, we used a Kto reporter line that expresses Kto fused with superfolder GFP (Kto-GFP) at the C-terminus under the control of its endogenous promotor ^51^ and an anti-Kto antibody ^5^. Both the Kto-GFP reporter and Kto immunostaining showed that Kto was ubiquitously expressed in both type I and type II NBs and all their progeny (Figure 2A-B’). Furthermore, the endogenous Kto expression in type II NB lineages was largely abolished by the expression of *UAS-Kto RNAi* driven by *pntP1-GAL4* (Figure 2C-C’), supporting that the Kto knockdown phenotype is indeed caused by the loss of Kto. For examining Skd expression, we used *UAS-myr-Tdtom* reporter expression driven by *skd-GAL4*, in which GAL4 is expressed under the control of an *skd* enhancer element ^52^. The reporter showed that *skd-GAL4* is expressed in both type I and type II NBs and their progeny in *Drosophila* larval brain, with relatively stronger expression in NBs and their young progeny close to the NBs (Figure 2D-D”), suggesting that Skd is also ubiquitously expressed but more enriched in NBs and newly generated progeny. Similar ubiquitous expression patterns of Kto and Skd also support that they could function together.

**Figure 2.**
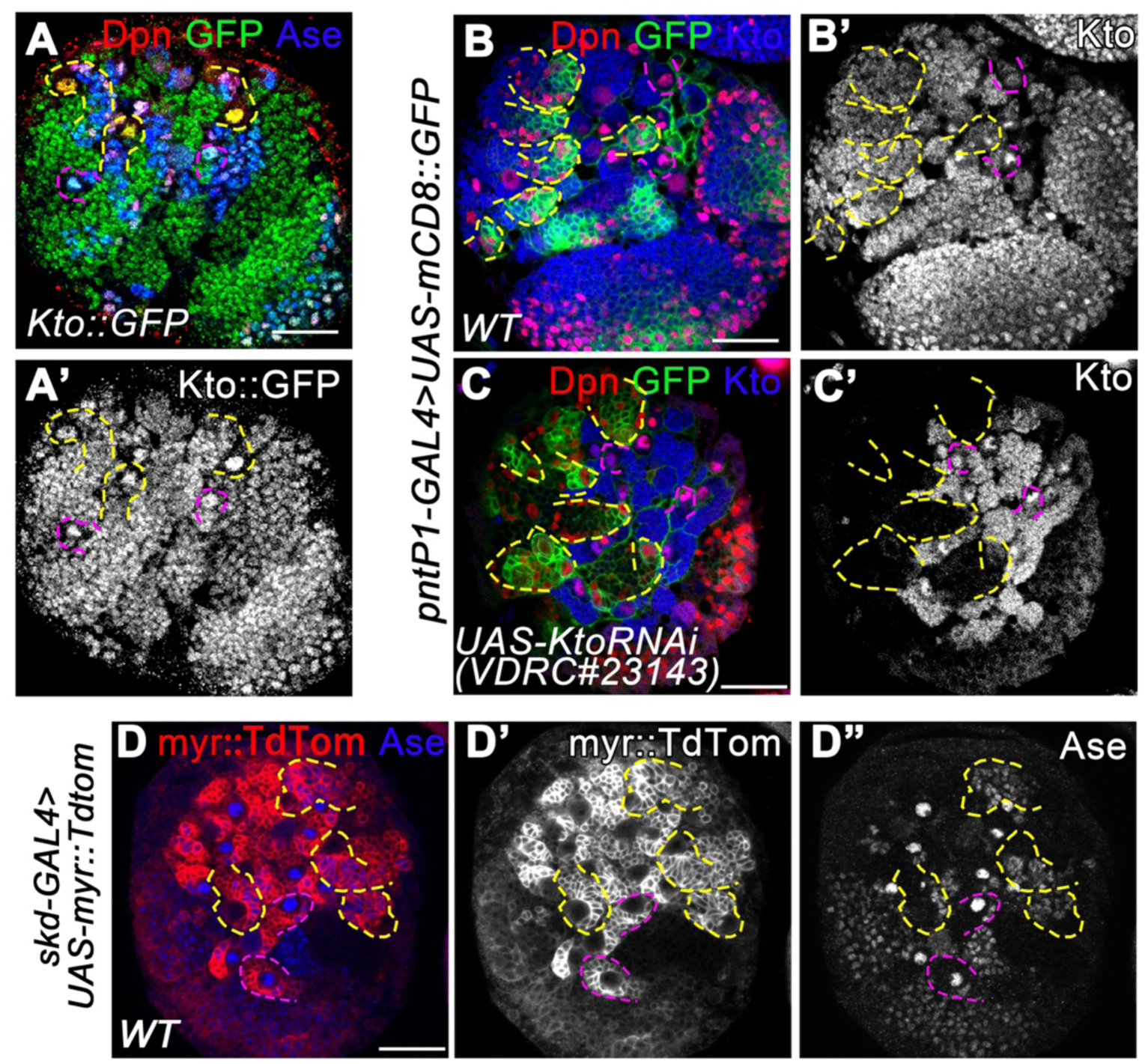
Skd and Kto are ubiquitously expressed in *Drosophila* larval brains. (A-A’) Kto-GFP is expressed in both type I (outlined with purple dashed lines) and type II (yellow dashed lines) NB lineages. Type II NB lineages are identified by the absence of Ase in the NB and the presence of multiple Dpn^+^ Ase^+^ INPs, whereas type I NBs are identified by the presence of Ase. (B-C’) Endogenous Kto, detected by anti-Kto antibody staining, is expressed in both type I (e.g. purple dashed lines) and type II (labeled with mCD8-GFP driven by *pntP1-GAL4*, yellow dashed lines) in a wild type brain (B-B’). Expression of *UAS-Kto RNAi* driven by *pntP1-GAL4* largely abolished endogenous Kto expression in type II NB lineages (C-C’). (D-D”) Myr::tdTom driven by *skd-GAL4* is expressed in all type II NB (Ase^-^) lineages (yellow dashed lines) and type I NB (Ase^+^) lineages (e.g. purple dashed lines). Scale bars equal 50µm.

### Knockdown of Skd/Kto leads to loss of Erm expression and persistence of PntP1 expression in INPs

We next tried to determine if the supernumerary type II NBs resulting from Skd and Kto knockdown were generated from dedifferentiation of imINPs. Previous studies have shown that Erm and Six4 are required to prevent dedifferentiation of imINPs by forming a complex with PntP1 to inhibit PntP1 activity and expression ^26,41,42,53^. Therefore, we examined Erm and Six4 expression after knocking down Skd/Kto. In wild type larval brains, Six4 is expressed in type II NBs and their progeny including imINPs and mINPs, whereas Erm is expressed in imINPs (except the newly born one) ^26,42^ (Figure S1A-A”, Figure 3A-A”). We found that knockdown of Skd or/and Kto in type II NB lineages did not obviously affect the expression levels of Six4-GFP (Figure S1B-D”). However, when either Skd or Kto was knocked down, Erm levels were reduced by 70% (Figure 3B-D). Consistently, the expression of PntP1 in INPs failed to be repressed when Skd or Kto was knocked down. In normal type II NB lineages, PntP1 is normally expressed in 3-4 imINPs but not in mINPs labeled with E(spl)mγ-GFP ^34^ (Figure 3E-E”, H), whereas in Skd or Kto knockdown lineages, the number of PntP1^+^ INPs almost doubled and PntP1 was also detected in E(spl)mγ-GFP^+^ mINPs (Figure 3F-H), indicating that the suppression of PntP1 expression in mINPs was delayed due to the loss of Erm expression. These results suggest that Skd and Kto are required for the activation of Erm expression in imINPs.

**Figure 3.**
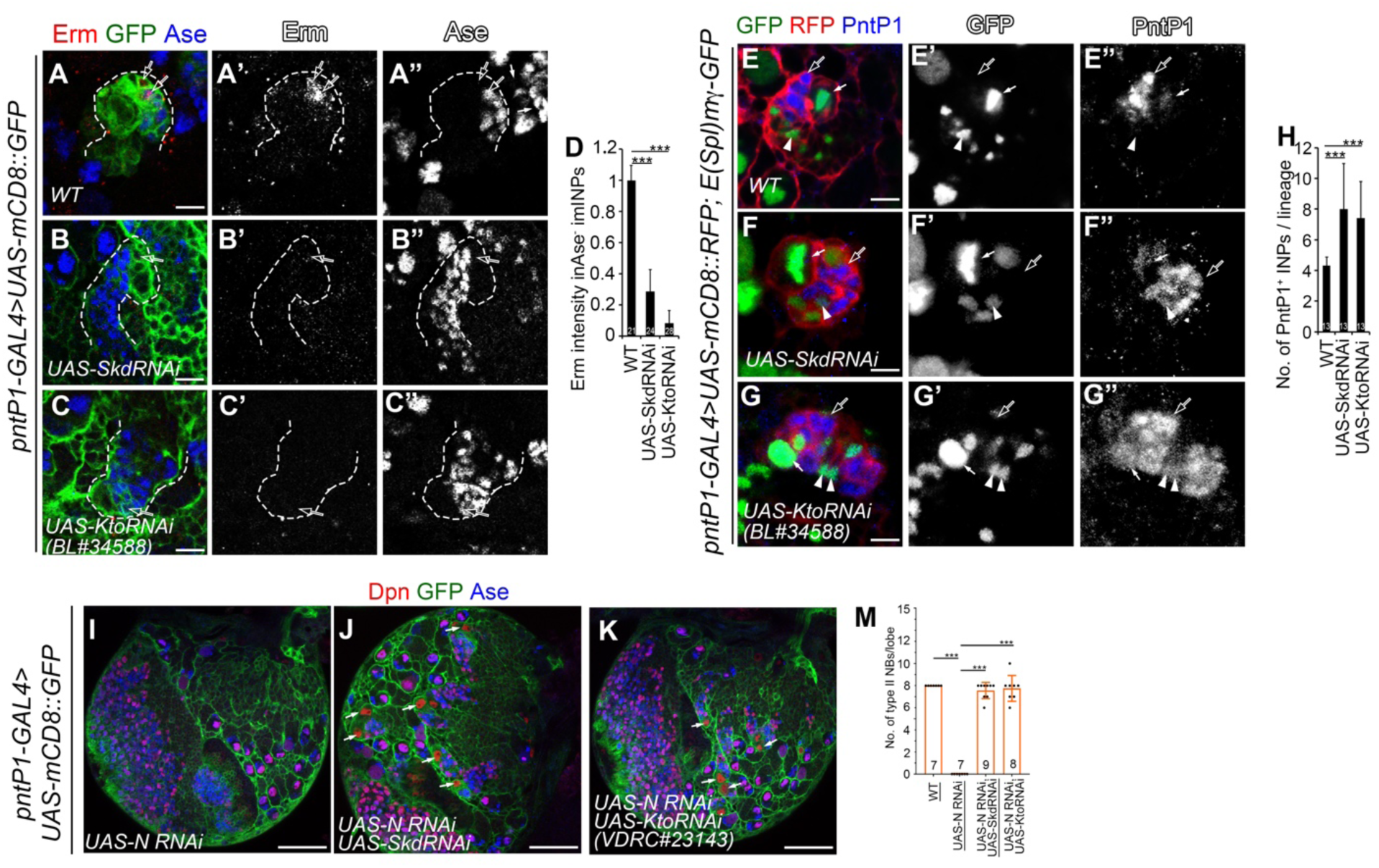
Skd and Kto are required for the activation of Erm expression. Type II NB lineages are labeled with mCD8-GFP (A-C”, I-K) or mCD8-RFP (E-G”) driven by *pntP1-GAL4* and counterstained with anti-Erm, anti-Ase, anti-PntP1, and/or anti-Dpn antibodies. Open arrows point to imINPs (A-C”, E-G”), arrowheads to E(Spl)mγ-GFP^+^ mINPs in (E-G”), and white solid arrows to type II NBs in (E-G”, I-K). Scale bars equal 10µm in (A-C”, and E-G”) or 50µm in (I-K). (A-A”) In a wild type type II NB lineage, Erm is expressed in both Ase^-^ and Ase^+^ imINPs. (B-D) Knockdown of Skd (B-B”) or Kto (C-C”) in type II NB lineages largely abolishes Erm expression in imINPs. Quantifications of the Erm staining intensity is shown in (D). Data represent Mean ± SD. ***, *P* < 0.001. The number on each bar represents the number of imINPs examined. (E-E”) PntP1 is expressed in type II NBs, imINPs, but never in E(spl)mγ^+^ mINPs in a wild type type II NB lineage. (F-H) Knockdown of Skd (F-F”) or Kto (G-G”) results in an increased number of PntP1^+^ cells and persistence of PntP1 expression in E(spl)mγ^+^ mINPs. Quantifications of the number of PntP1^+^ INPs are shown in (H). Data represent Mean ± SD. ***, *P* < 0.001. The number on each bar represents the number of type II NB lineages examined. (I-L) Knockdown of Notch eliminates all type II NBs in a third instar larval brain (I), but concomitant knockdown of Skd (J) or Kto (K) rescues the loss of type II NBs resulting from Notch knockdown. Quantifications of the number of type Ase^-^ type II NBs are shown in (L). Data represent Mean ± SD. ***, *P* < 0.001. The number on each bar represents the number of brain lobes examined.

To further confirm that Skd/Kto is required for Erm expression, we examined if the loss of type II NBs resulting from Notch knockdown could be rescued by Skd/Kto knockdown. Our previous studies show that the loss of type II NBs resulting from the loss of Notch is due to precocious activation of Erm by PntP1 in type II NBs ^41^. Indeed, our results showed that expressing either *UAS-Skd RNAi* or *UAS-Kto RNAi* in type II NBs fully rescued the loss of type II NB lineages resulting from Notch knockdown (Figure 3K-L), suggesting that the ectopic activation of Erm in Notch knockdown type II NBs also requires Skd and Kto.

### The supernumerary type II NBs resulting from Skd/Kto knockdown are generated through dedifferentiation of imINPs due to the loss of Erm expression

Next, we wanted to determine whether the supernumerary type II NBs resulting from Skd/Kto knockdown are indeed generated from dedifferentiation of imINPs due to the loss of Erm. To this end, we first tested if simultaneously knocked down Skd and Kto specific in imINPs could cause supernumerary type II NBs and if the phenotypes can be rescued by the expression of Erm. We knocked down Skd and Kto in imINPs with *erm-GAL4(II)* ^53,54^ in the presence of *E(spl)mγ-GFP.* Our results showed that double knockdown of Skd and Kto in imINPs increased the number of type II NBs to about 18 per lobe (Figure 4A-C, M). However, simultaneous expression of UAS-Erm in imINPs completely suppressed the generation of supernumerary type II NBs resulting from Skd and Kto knockdown (Figure 4D, M).

**Figure 4.**
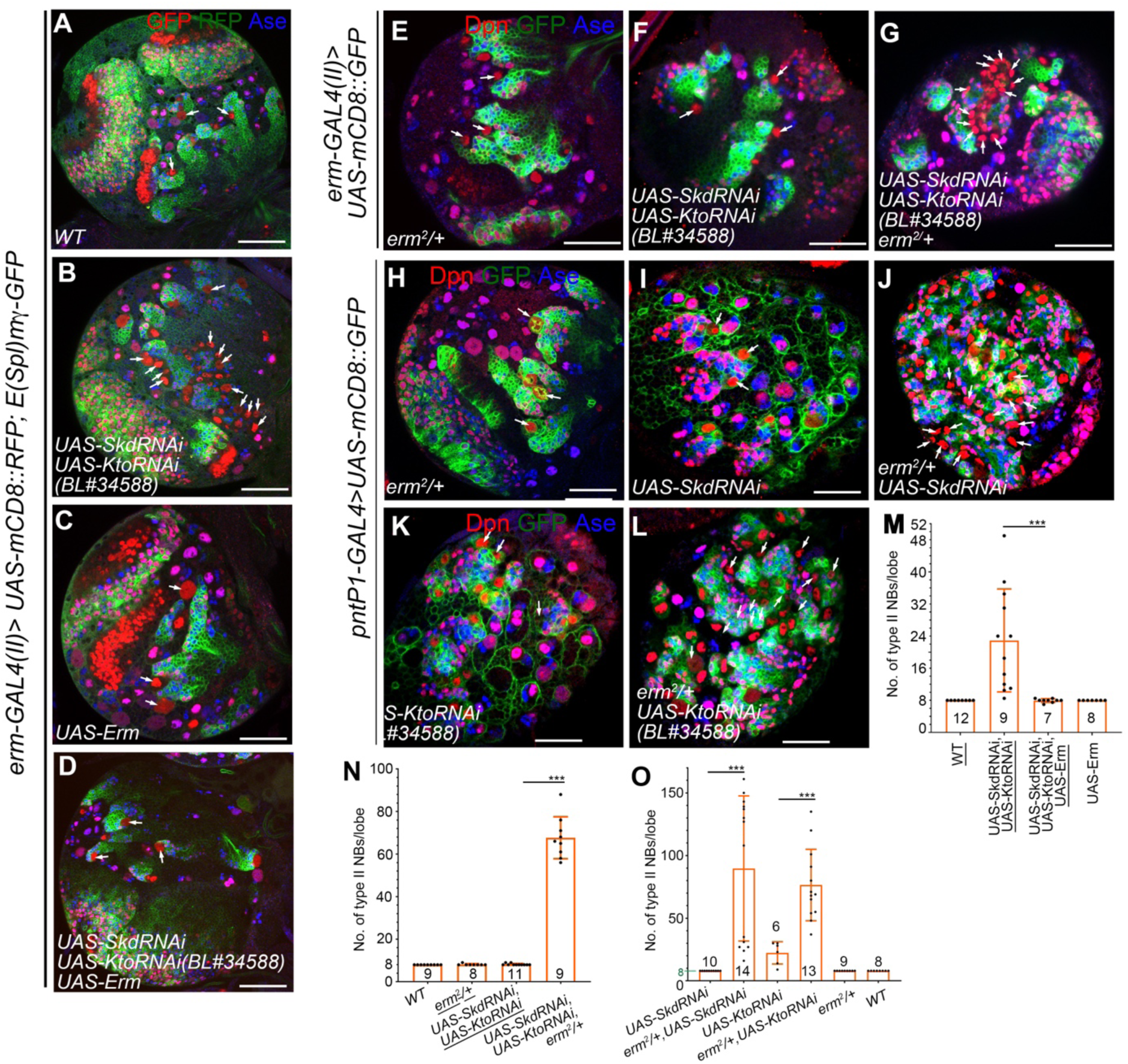
Supernumerary type II NBs resulting from Skd/Kto knockdown arise from dedifferentiation of imINPs due to the loss of Erm expression. In all images, type II NB lineages are labeled with mCD8-RFP (A-D) or mCD8-GFP (E-L) driven by *erm-GAL4(II)* (A-G) or *pntP1-GAL4* (H-L) and counterstained with anti-Dpn and/or Anti-Ase antibodies. White arrows point to type II NBs. Scale bars equal 50µm. (A) A wild type brain lobe with the *E(spl)mγ-GFP* transgene contains only eight type II NBs. (B-D, M) Double knockdown of Skd and Kto with *erm-GAL4(II)* in imINPs in the presence of *E(spl)mγ-GFP* leads to the generation of supernumerary type II NBs (B). Expression of *UAS-Erm* driven by *erm-GAL4(II)* does not affect the number of type II NBs (C), but rescues the supernumerary type II NB phenotype resulting from Skd and Kto knockdown (D). Quantifications of the number of type II NBs are shown in (M). Data represent Mean ± SD. ***, *P* < 0.001. The number on each bar represents the number of brain lobes examined. (E-G, N) No ectopic type II NBs are generated in an *erm^2^/+* heterozygous mutant brain (E) or when Skd and Kto are simultaneously knocked down in imINP with *erm-GAL4(II)* in the absence of *E(spl)mγ-GFP* (F). However, double knockdown of Skd and Kto in with *erm-GAL4(II)* in the *erm^2^/+* background induces supernumerary type II NBs (G). Quantifications of the number of type II NBs are shown in (N). Data represent Mean ± SD. ***, *P* < 0.001. The number on each bar represents the number of brain lobes examined. (H-L, O) No extra type II NBs are generated in the *erm^2^/+* heterozygous mutant brain (H). Knockdown of Skd or Kto with *pntP1-GAL4* Knockdown in the wild type background gives rise no or weak supernumerary type II NB phenotypes, respectively (I, K). However, knockdown of Skd or Kto in the *erm^2^/+* heterozygous mutant background leads to synergistic enhancement of supernumerary type II NB phenotypes (J, L). Quantifications of the number of type II NBs are shown in (O). Data represent Mean ± SD. ***, *P* < 0.001. The number on each bar represents the number of brain lobes examined.

To further confirm that the loss of Erm is responsible for the generation of supernumerary type II NBs resulting from Skd/Kto knockdown, we tested genetic interactions between Erm and Skd/Kto by examining if the supernumerary type II NB phenotype resulting from Skd/Kto knockdown would be enhanced in the *erm^2^/+* heterozygous mutant background in the absence of E(Spl)mγ-GFP. Indeed, while double knockdown of Skd and Kto with *erm-GAL4(II)* or single knockdown of Skd or Kto with *pntP1-GAL4* in the wild type background produced no or only weak supernumerary type II NB phenotypes (Figure 4F, I, K, N, O), their phenotypes were synergistically enhanced in *erm^2^/+* heterozygous mutant background (Figure 4E, G, H, J, L, N, O). Therefore, Skd and Kto genetically interact with Erm in preventing dedifferentiation of imINPs. Taken together, these results demonstrated that Skd and Kto are required to prevent dedifferentiation of imINPs by mediating the activation of Erm in imINPs.

### Skd and Kto are essential cofactors of PntP1

The mediator complex has been shown to regulate target gene expression by directly or indirectly interacting with sequence-specific transcription factors ^1,2^. Since Erm expression in imINPs is activated by PntP1 ^25,34,41^, we next investigated whether Skd and Kto are required for PntP1’s function. Our previous studies have shown that misexpression of PntP1 in type I NBs suppresses Ase expression and induces the generation of INP-like cells ^34,38^. Since Skd and Kto are also expressed in type I NBs, we assessed the necessity of their functions in the transformation from type I NBs into type II-like NBs induced by PntP1. We focused on larval ventral nerve cords (VNCs), which contain only type I NB lineages, for phenotypic analyses. Our results showed that more than 90% of type I NBs lost Ase expression and about 15% of them generated Dpn^+^ Ase^+^ mINP-like cells when PntP1 was misexpressed in type I NBs (Figure 5A-B, E-F). Knockdown of Skd and Kto did not affect the expression of Ase in type I NBs (Figure 5C, E-F). However, when Skd and Kto were simultaneously knocked down, misexpressing PntP1 could suppress Ase in only 10% of type I NBs and failed to induce any INP-like cells (Figure 5D-F), indicating that Skd and Kto are essential for PntP1’s function.

**Figure 5.**
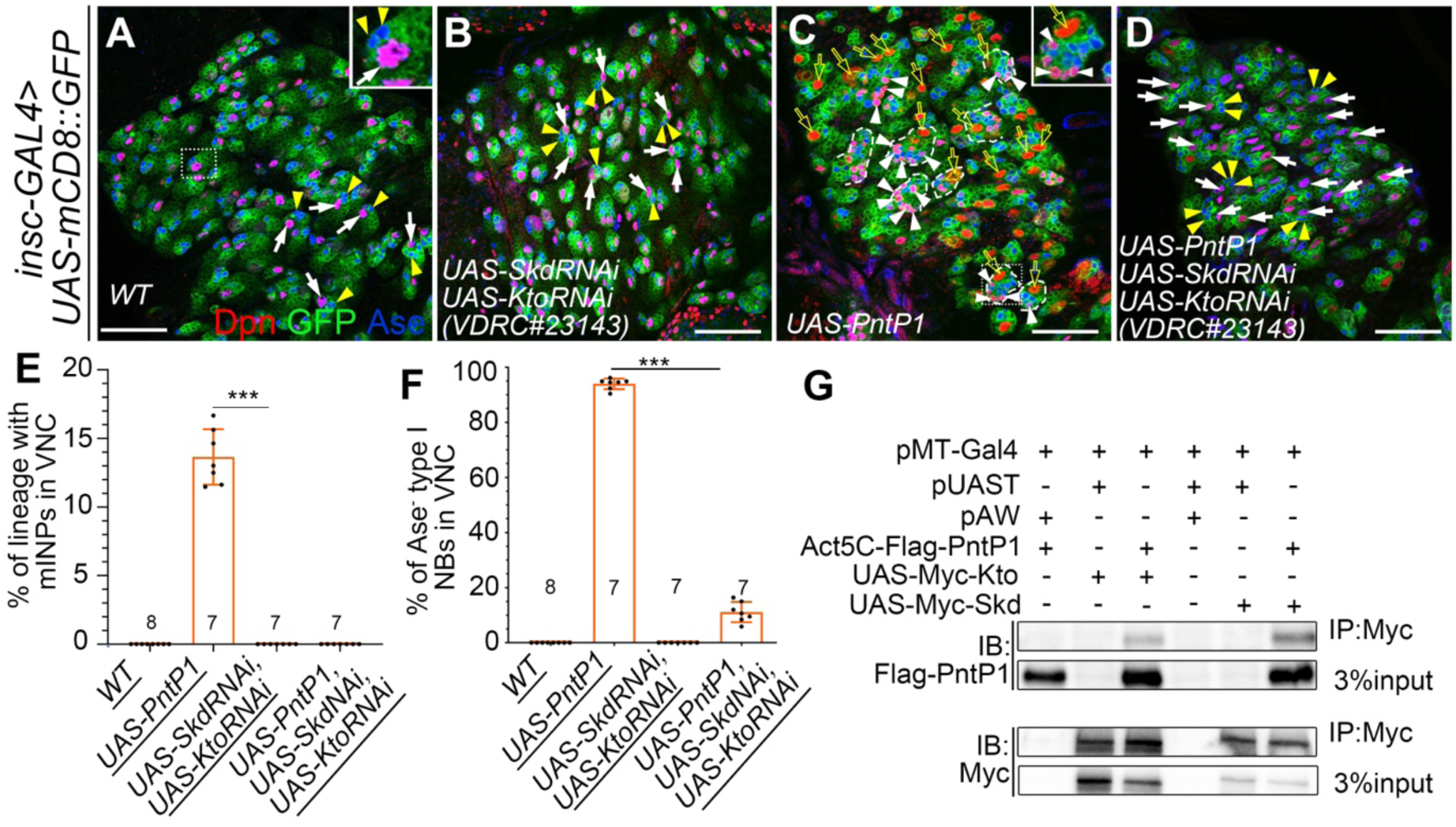
Skd and Kto are required for PntP1 to suppress Ase expression and induce INP-like cells in type II NB lineages. In all images, type I NB lineages in the VNC are labeled with mCD8-GFP driven by *insc-GAL4* and counterstained with anti-Dpn and anti-Ase antibodies. Scale bars equal 50µm. (A-B) All type I NBs express Ase (white arrows) and directly generated GMCs (small Ase^+^ Dpn^-^ cells, yellow arrowheads) in the wild type (A) and Skd and Kto double knockdown brains (B). The inset in (A) is an enlarged view of the lineage within the dotted square. (C) Misexpressing PntP1 in type I NBs represses Ase expression in the majority of type I NBs (open arrows) and induces Ase^+^ Dpn^+^ mINP-like cells (arrowheads) in a subset of lineages (dashed lines) (C). The inset shows an enlarged view of the lineage highlighted with a dotted square with mINP-like cells. (D) All type I NBs remain Ase^+^ and generate GMCs (yellow arrowheads) directly when Skd and Kto are simultaneously knocked down while PntP1 is misexpressed in type I NBs. (E-F) Quantifications of the percentage of type I NB lineage with mINP-like cells (E) and the percentage of Ase^-^ type I NBs (F) in VNCs with indicated genotypes. Data represent Mean ± SD. ***, *P* < 0.001. The number on each bar in both (E) and (F) represents the number of VNCs examined. (G) Biochemical interactions between Flag-PntP1 and Myc-Kto or Myc-Skd are detected in S2 cells transfected with their corresponding expression constructs by co-IP and western blotting analyses.

To investigate whether Skd/Kto form a complex with PntP1, we performed co-immunoprecipitation (co-IP) followed by western blot analysis. We co-expressed Flag-tagged PntP1 and Myc-tagged Skd or Myc-tagged Kto in *Drosophila* S2 cells. Our results showed that immunoprecipitation of PntP1 with an anti-Flag antibody resulted in co-precipitation of Skd or Kto (Figure 5G), demonstrating that Skd and Kto form a complex with PntP1. We were also able to co-IP Skd and Kto when Myc-tagged Skd and HA-tagged Kto were co-expressed in S2 cells (data not shown) as shown in previous studies ^5^. Thus, the biochemical results demonstrate that PntP1 forms a complex with Skd and Kto and that Skd/Kto are indispensable for PntP1 to activate Erm expression and suppress Ase expression.

Our previous studies report that during maturation of imINPs, Erm together with Six4 inhibit PntP1 activity by forming a complex with PntP1 ^42^. Since here we demonstrate that PntP1 activity also requires formation of a complex with Skd and Kto, we asked if Six4 and Erm inhibit PntP1 activity by displacing Skd/Kto from the PntP1 complex or simply by binding to the PntP1-Skd/Kto complex. Therefore, we also examined if Six4 also forms a complex with Skd and Kto by performing co-IP in S2 cells. Our results show that when Six4-HA was co-expressed with myc-Skd or myc-Kto in S2 cells, Six4 could be co-IPed with Skd or Kto (Figure S2). These results, together with our previous results that Six4, Erm, and PntP1 could be co-IPed in S2 cells ^42^, suggest that Six4 and Erm likely inhibit PntP1 activity by forming a complex with PntP1, Skd, and Kto rather than displacing Skd/Kto from the complex.

### Skd and Kto likely function independently of the kinase module subunits CDK8 and CycC in preventing the generation of supernumerary type II NBs

Since Skd and Kto may or may not form a subcomplex with the other two mediator kinase module subunits CDK8 and CycC to regulate target gene transcription ^55^, we next tested if CDK8 and CycC could be involved in preventing the generation of supernumerary type II NBs. To this end, we either knockdown CDK8 or CycC alone or together with Skd in type II NB lineages using *pntP1-GAL4* as a driver. Simultaneous knockdown of CDK8 or CycC together with Skd would allow us to test if they genetically interact. Furthermore, knockdown Skd could provide a sensitized background to detect CDK8 or CycC knockdown phenotypes. We performed these knockdown experiments in the absence of *E(spl)mγ-GFP* to make it easier to detect synergistic enhancement of the phenotypes if any. Our results show that knockdown of CDK8 or CycC alone or together with Skd didn’t cause generation of any extra type II NBs (Figure 6A-C, M-O, Y). These results suggest that Skd and Kto may not require CDK8 and CycC to prevent dedifferentiation of imINPs.

**Figure 6.**
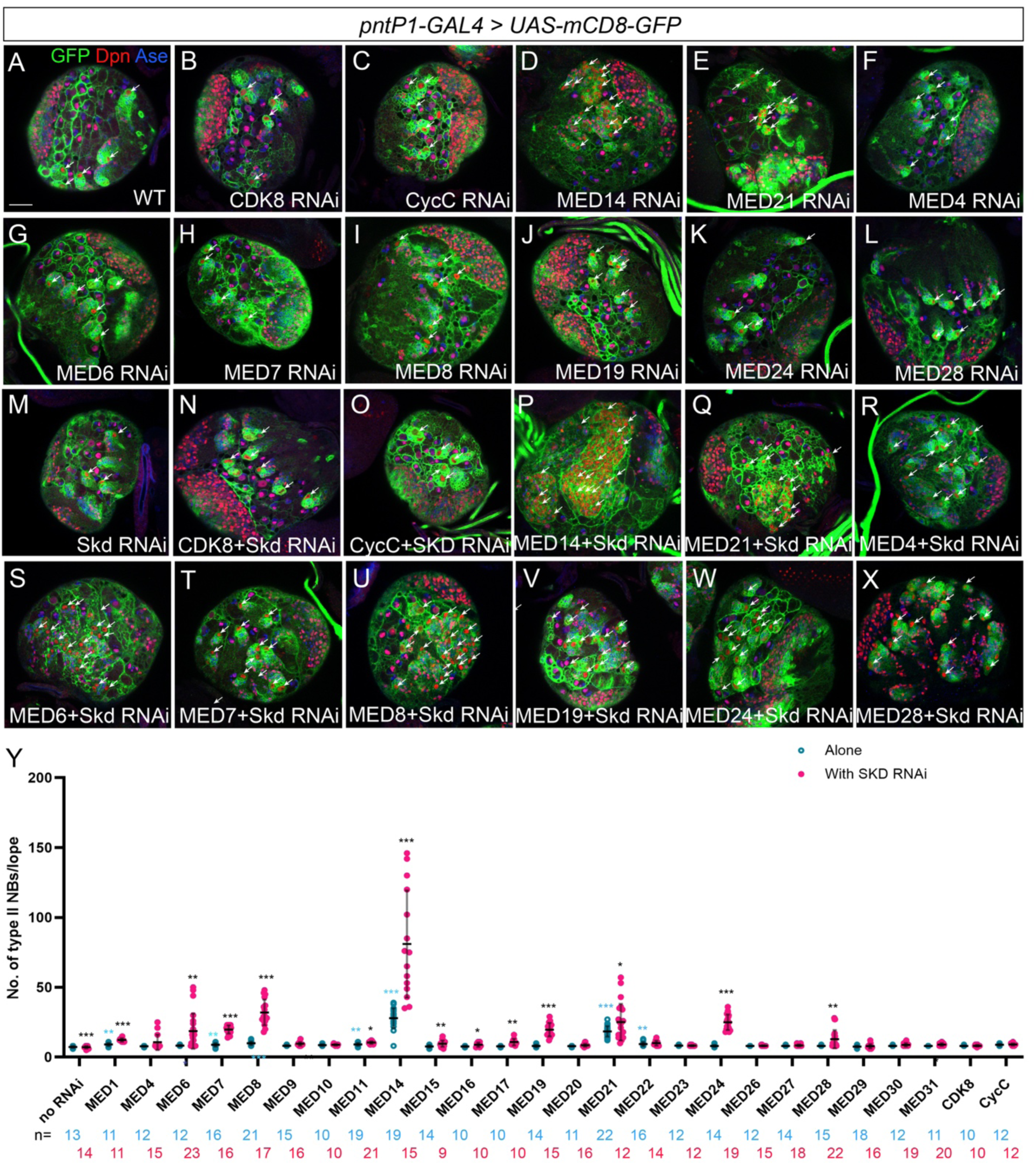
A subset of the core mediator complex subunits genetically interact with Skd in preventing the generation of supernumerary type II NBs. In all images, type II NB lineages are labeled with mCD8-GFP driven by *pntP1-GAL4* and counterstained with anti-Ase and Anti-Dpn antibodies. Arrows point to type II NBs. Scale bar: 50µm. (A) A wild type larval brain lobe consistently contains eight type II NBs. (B-L) show larval brain lobes with knockdown of CDK8 (B), CycC (C), or subunits of the mediator core complex (D-L) in the wild type background. (M-X) show brain lobes with Skd knockdown alone (M) or knockdown of Skd together with CDK8 (N), CycC(O), or the mediator core complex subunits (P-X). (Y) Quantifications of number of type II NBs in brains with indicated genotypes. Data represent Mean ± SD. ***, *P* < 0.001. **, *P* < 0.01. *, *P* <0.05. Asterisks in black indicate statistical significance compared with knockdown of the same Mediator subunit alone, whereas asterisks in cyan indicate statistical significance compared with the wild type without RNAi. The sample size (n) for each group represents the number of brain lobes examined.

### A specific subset of the mediator core complex components likely function cooperatively with Skd and Kto to prevent supernumerary type II NBs

Skd and Kto can function through the core mediator complex to regulate target gene expression or just function as a submodule independently of the core complex ^5^. Therefore, we asked if components of the core mediator complex could be involved in preventing the generation of supernumerary type II NBs. We performed similar genetic interaction tests with Skd for each of the 26 known subunits of the core mediator complex as CDK8 and CycC. Our results showed that when these subunits were knocked down alone, only MED14 or MED21 gave obvious supernumerary type II NB phenotypes (about total 20 or 35 type II NBs per lobe on average, respectively) (Figure 6 D, E, Y), others generated no or less than 1 extra type II NBs per brain lobe on average (Figure 6F-L, Y, and data now shown). However, when Skd was knocked down simultaneously, knockdown of MED6, MED7, MED8, MED14, MED19, MED21, and MED24, consistently led to synergistic enhancement of the supernumerary type II NB phenotype, with an average number of type II NBs ranging from 17 to 72 per lobe (Figure 6P,Q, S-W, Y), whereas knockdown of other mediator core complex subunits gave no or only less than 5 (mostly just 1-2) extra type II NBs (Figure 6R, X, Y). Among all the mediator core complex subunits, knockdown of MED14 gave the strongest enhancement of the phenotype, which is consistent with its critical role as a central backbone that links all three main submodules (head, middle and tail) of the mediator core complex and maintains the proper architecture and function of the complex ^2^. These genetic interaction data suggest that Skd and Kto could be function together with a specific subset of the mediator core complex subunits to prevent the generation of supernumerary type II NBs.

### Skd and Kto are required to maintain self-renewal of type I NBs

Since Skd and Kto are ubiquitously expressed not only in type II NB lineages but also also in type I NB lineages in larval brains, we then asked if Skd and Kto have any role in the development of type I NB lineages. Previous studies have shown that overexpression of Kto prevents NB shrinkage and termination of self-renewal at early pupal stages possibly by antagonizing the function of the core mediator complex and preventing the core mediator complex and ecdysone receptor (EcR) from activating metabolic genes, but the function of Skd and Kto in larval type I NBs has never been investigated. Therefore, we simultaneously knocked down Skd and Kto in type I NBs using a type I NB-specific driver *ase-GAL4* ^23,24^ or a pan-NB driver *insc-GAL4* ^56^ and examined the number of type I NBs at different larval stages. Our results showed that knockdown of Skd and Kto led to a decrease in the number of type I NBs in the central brain (Figure 7A1-B3, E) and the VNC (Figure 7H-I, K) at 4 days after egg lying (AEL) by approximately 20% and 15%, respectively, but not at earlier stages. Further, the diameter of Skd and Kto double knockdown type I NBs and the number of their associated GMCs were also significantly decreased (Figure 7F-G) at 3-4 days AEL. The reduced number of GMCs was likely due to a decreased mitotic rate of type I NBs as the percentage of pH3^+^ type I NBs was decreased by 50% (Figure S3A-C). These results indicate that Skd and Kto are also required to maintain the self-renewal and proliferation of type I NBs during mid-late larval stages.

**Figure 7.**
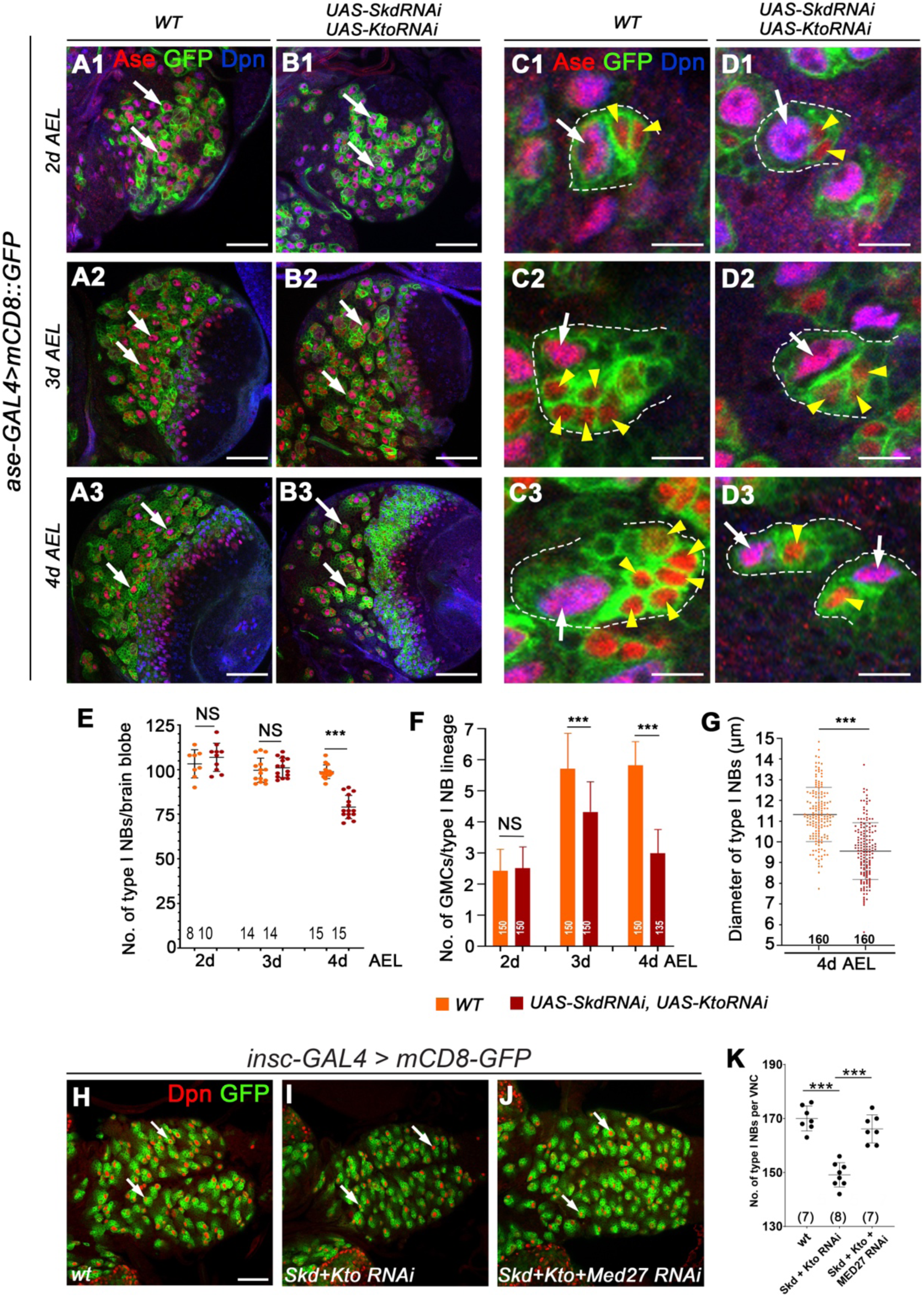
Double knockdown of Skd and Kto decreases the number and size of type I NBs and the number of associated GMCs. In all images, type I NB lineages were labelled with mCD8-GFP driven by *ase-GAL4* (A1-D3) or *insc-GAL4* (H-J) and counterstained with anti-Ase and anti-Dpn antibodies. Arrows point to type I NBs. Scale bars equal 50µm in (A1-B3, H-I) or 10µm in (C1-D3). (A1-B3) Wild type (A1-A3) brain lobes and brains lobes with Skd Kto double knockdown in type I NBs at 2 - 4 days egg laying (AEL). (C1-D3) Enlarged views of a single wild type (C1-C3) and Skd Kto double knockdown (D1-D3) type I NB lineages (outlined with dashed lines) at 2 - 4 days AEL. Yellow arrowheads point to GMCs. (E-G) Quantifications of the number of type I NBs (E) and GMCs (F), and the diameter of type I NBs (G) in wild type and Skd Kto double knockdown brain lobes at 2 - 4 days AEL. Data represent Mean ± SD. ***, *p* < 0.001; NS, no significance, student *t*-test. The number on each bar represents the number of brain lobe (E), the number of type I NB lineages (F), or the number of type I NBs (F). (H-K) A wild type VNC (H) and VNCs with Skd Kto double knockdown (I) or Skd Kto Med27 triple knockdown with insc-GAL4. Quantifications of the number of type I NBs are shown in (K). Data represent Mean ± SD. ***, *p* < 0.001, student *t*-test. The number on each bar represents the number of VNCs examined.

To test if Skd and Kto maintain the NB self-renewal by antagonizing the function of the mediator core complex and EcR, we then examine if inhibiting EcR or knocking down mediator core complex subunits that have been shown to promote NB shrinkage at early pupal stages would rescue the loss of type I NBs. Indeed, we found that knocking down Med27 or expressing dominant negative form of EcR (EcR^DN^) largely rescued the loss of type I NBs caused by Skd Kto double knockdown (Figure 7J-K, Figure S3D-G), indicating that the loss of type I NBs caused by Skd and Kto knockdown is due to de-inhibition of the activity of the core mediator complex and EcR.

## DISCUSSION

In this study, we revealed two novel functions of Skd and Kto in *Drosophila* larval NB lineages. One is to prevent the generation of tumorigenic supernumerary type II NBs by promoting the maturation of imINPs. Another is to maintain the self-renewal of NBs during larval stages. Skd and Kto perform these two functions likely by functioning through two distinct subsets of the mediator core complex subunits to regulate different gene expression (Figure S4).

In type II NB lineages, the primary Skd/Kto knockdown phenotype is the supernumerary type II NBs. We demonstrate that Skd and Kto prevents the generation of supernumerary type II NBs by forming a complex with PntP1 to mediate the activation of Erm by PntP1. In addition to activating Erm, PntP1 also suppresses Ase expression by activating Tailless (Tll) ^35–37^. In type I NBs, we demonstrate that suppression of Ase by mis-expressed PntP1 also requires Skd and Kto. Concomitant knockdown of Skd/Kto prevented the suppression of Ase in type I NBs by mis-expressed PntP1. However, knockdown of Skd/Kto in type II NB lineages with *pntP1-GAL4* did not lead to ectopic activation of Ase in type II NBs. One reason could be that *pntP1-GAL4* has relatively weak expression in type II NBs compared to its expression in imINPs. Therefore, expression of *UAS-Skd/Kto RNAi* driven by *pntP1-GAL4* may not fully eliminate Skd or Kto from type II NBs and residual expression of Skd or Kto may persist at below detectable levels. Another reason, which we think is more likely, could be that other factors expressed in type II NBs, such as Btd, may also function to suppress Ase. Although loss of Btd in type II NB lineages mainly leads premature differentiation of INPs into GMCs due to ectopic activation of Pros expression in newly generated imINPs, misexpression of Btd also suppresses Ase expression in a small percentage of type I NBs and co-expression of Btd and PntP1 can transform nearly all type I NB lineages into type II NB lineages ^38,39^. Therefore, PntP1 and Btd may have partial redundant role in suppressing Ase although PntP1 plays a dominant role. Knockdown of Skd/Kto may only impact the function of PntP1 but not Btd. Thus, Ase remains suppressed in Skd/Kto knockdown type II NBs. The persistence of Ase expression likely activates Prospero expression ^57,58^, which promotes the progeny to adopt the fate of GMC and exit cell cycle ^59^ and prevent them from dedifferentiating back into NBs in the absence of Skd and/Kto. Our results that Skd and Kto only mediate the suppression of Ase by PntP1 but not by Btd also indicates that although Skd and Kto is ubiquitously expressed, Skd and Kto may not generally mediate gene activation by any transcription factors even in the same cell. Rather, they may only interact with specific transcription factors in any given cell.

Skd and Kto can form the kinase module together with CycC and CDK8 and modulate the activity of the medicator core complex but can also form a submodule without CycC and CDK8 and regulate target gene expression independently of CycC and CDK or the core complex ^60^. Our results that knockdown of CycC or CDK8 neither caused the supernumerary type II NB phenotype nor synergically enhanced the supernumerary type II NB phenotype resulting from Skd knockdown argue against the involvement of CycC and CDK8 in the regulation of INP maturation by Skd and Kto. However, Skd and Kto likely still function together with a subset of mediator core complex subunits to prevent the generation of super numerary type II NBs, particularly MED6, MED7, MED8, MED14, MED19, MED21, and MED24, as knockdown of these subunits consistently caused supernumerary type II NB phenotypes or/and synergistically enhanced the supernumerary type II NB phenotypes when Skd was simultaneously knockdown. Previous studies only show that the mediator core complex components are required for NB shrinkage and termination during early pupal stages ^30,32^. Thus, our work reveals a novel function of the mediator core complex in preventing the generation of supernumerary type II NBs.

In addition to preventing dedifferention of imINPs, our results also show that Skd and Kto are required to maintain the self-renewal of type I NBs at late larval stages by antagonizing the function of the mediator core complex and EcR. Previous studies reported that the mediator core complex promotes NB shrinkage and termination of self-renewal only at early pupal stages by mediating the activation of metabolic genes by ecdysone receptors ^30,32^. Findings from this work suggest that the mediator complex and EcR could also function at larval stages to inhibit NB self-renewal and proliferation, but their functions are normally inhibited by Skd and Kto. Such inhibitory effects of Kto on the mediator core complex has also been shown my manipulating Kto expression at early pupal stages ^30,32^. However, during early pupal stages, possible downregulation of Kto ^32^ and a large ecdysone peak ^61^ allow the mediator core complex and EcR to activate metabolic genes to terminate NB self-renewal.

The subunits of the core mediator complex that are required for termination of NB self-renewal at early pupal stages include Med4, Med6, Med9, Med10, Med11, Med22, Med27, and Med31 ^30^, which are largely different from those required for preventing the generation of supernumerary type II NBs shown in this study, including Med6, Med7, Med8, Med14, Med19, Med21, and Med24. Therefore, the mediator core complex subunits can be divided into two distinct subsets. One subset functions together with Skd and Kto to promote type II NB specification and INP maturation, whereas the other subset is normally inhibited by Skd and Kto during NB self-renewal at larval stages. How can Skd and Kto coordinate with one subset of the core mediator complex subunits but antagonize the function of another subset? One possibility could be that these two different subsets of core mediator complex subunits exist in two separate smaller core complexes. Binding of Skd and Kto to one subset activates their transcriptional activity or promote the binding of sequence-specific transcription factor such as PntP1 to the core mediator complex. In contrast, binding of Skd and Kto to the other subset inhibits the transcriptional activity of the core complex or interfere the binding of sequence specific transcription factor (such as ecdysone receptor) to the core mediator complex. Alternatively, these two distinct subsets of the core mediator complex subunits still coexist in the same complex. However, binding of Skd and Kto may cause conformational changes of the core mediator complex that favors the binding of PntP1 to the core complex but disrupts the binding of ecdysone receptor to the complex. It has been shown that different sequence-specific transcription factors bind to different subunits in the core mediator complex to activate gene transcription ^62,63^. To distinguish these possibilities will require proteomics and structural analyses of the core mediator complex in the *Drosophila* NBs in the future.

## METHODS

### Fly stocks

The *E(spl)mγ-GFP* reporter line ^64^ was used to label NBs and mINPs. The *Kto-GFP* line (#318680, Vienna Drosophila Resource Center [VDRC], Vienna, Austria) was used to detect Kto expression. The *Six4-GFP* line (#67733, Bloomington *Drosophila* Stock Center [BDSC]) was used to detect Six4 expression. GAL4 lines, including *pntP1-GAL4* (also called *GAL4*^14–94^) ^34^, *erm-GAL4(II)* ^54,65^, *insc-GAL4* ^56^, *Skd-GAL4* ^52^ (#76783, BDSC) and *ase-*GAL4 ^23,24^ were used for *UAS-transgenes* expression. Knockdown Notch was carried out with *UAS-N RNAi* (#7078, BDSC). *UAS-Skd RNAi* (#34630, BDSC) and *UAS-Kto RNAi* (#34588, BDSC; #23142, VDRC; # 23143, VDRC) were used for knocking down Skd or Kto respectively. *UAS-EcR^DN^* (#9452, BDSC) was used for inhibiting EcR. *UAS-Skd.J* (#63800, BDSC) and *UAS-Kto.J* (#63801, BDSC) ^5^ were used to rescue Skd or Kto knockdown phenotypes. *UAS-PntP1* ^34^ was used for mis-expression of PntP1 in type I NBs. *erm^2^* ^26^ allele was used to reduce Erm expression. Other *UAS-RNAi* lines for knocking down mediator complex subunits are listed in Supplemental Table S1.

### UAS-transgene expression

For *RNAi* knockdown and overexpression of genes in type II NB lineages, embryos were collected for 4∼10hrs at 25°C and shifted to 30°C to boost the efficiency. *UAS-dcr2* was coexpressed with *UAS-RNAi* transgenes to enhance the RNAi knockdown efficiency. For PntP1 misexpression in type I NB lineages, *insc-Gal4* together with the temperature sensitive *tub-Gal80^ts^* ^66^ (#7019, BDSC) were used to drive the expression of *UAS-pntP1*. Embryos were collected for 8∼10hrs and raised to hatch at 18°C, then larvae were shifted to 30°C for 3∼4 days to maximize the efficiency.

### S2 cell culture and co-immunoprecipitation

pAFW-PntP1 (i.e., *Act5C-Flag-PntP1*) vector (gifts from Dr. H. Y. Wang) ^49^ was used for expressing Flag-PntP1 in S2 cells under the control of the *Act5C* promoter. pUAST-Myc-Skd (i.e., *UAS-Myc-Skd*), pUAST-Myc-Kto (i.e., *UAS-Myc-Kto*) and pUAST-HA-Kto (i.e., *UAS-HA-Kto*) (gifts from Dr. Y. Zhao) ^55^ were used to express Skd or Kto in S2 cells. Mock vectors pUAST and pAW (generated by deleting *Flag-PntP1* cassette from pAFW-PntP1) were co-transfected as controls to balance the amounts of each vector transfected. S2 cell culture and co-IPs flowed by western blotting were performed as described previously ^42^.

### Immunostaining and confocal microscopy

Brain lobes along with VNCs dissected from larvae at desired stages were immunostained as previously described ^31^ except that the fixation time was increased to 35 mins. Primary antibodies used are chicken anti-GFP (Catalog #GFP-1020, Aves Labs, Tigard, Oregon; 1:500-1000), rabbit anti-DesRed (Catalog #632392, Takara Bio USA, Inc., Mountain View, California; 1:250), rat anti-mCD8 (Catalog #12-0088-42, Thermo Fisher Scientific, Waltham, Massachusetts, 1:100), rabbit anti-Dpn (1:500) (a gift from Dr. Y.N. Jan) ^67^, guinea pig anti-Ase (a gift from Dr. Y.N. Jan, 1:5000) ^23^, rat anti-Erm (a gift of Dr. C. Desplan; 1:300), rabbit anti-PntP1 (a gift of Dr. J. B. Skeath; 1:500) ^68^, guinea pig anti-Kto (a gift from Dr. J.E. Treisman) ^5^, mouse anti-PH3 (Catalog # ab80612, Cambridge, Massachusetts, 1:500). Secondary antibodies conjugated to Daylight 405 (1:300-400), Daylight 488 (1:100), Cy3 (1:500), Rhodamine Red-X (1:500), Daylight 647 (1:500) or Cy5 (1:500) used for immunostaining are from Jackson ImmunoResearch (West Grove, Pennsylvania). Images were collected using a Carl Zeiss LSM780 confocal microscopy and processed with Adobe Photoshop.

### Statistics

Statistics analyses were performed with excel. Two-tailed unpaired student’s t-test was used for statistical analyses. Data represent Mean ± SD in all figures. Statistical significance was defined as p-value <0.05. * indicates *p*<0.05; **, *p*<0.01; and ***, p<0.001.

## Resource availability

## Materials availability

Fly lines and plasmids used in this study are available from the corresponding author without restriction.

## Data and code availability

This paper does not report proteome or transcriptome data or original code. Additional raw files and information required to reanalyze the data are available from the corresponding authro upon request.

## ACKNOWLEDGEMENTS

We thank Drs. Y. N. Jan, J. B. Skeath, H. Y. Wang, C. Desplan and Y. Zhao, J.E. Treisman, the Bloomington *Drosophila* Stock Center, Vienna *Drosophila* Resource Center, Zurich ORFeome Project, and *Drosophila* Genomics Resource Center for antibodies, plasmids and fly stocks; Drs. F. Pignoni and A. S. Viczian for sharing research facility; N Jusic for technique support; Drs. R. T. Matthews, F. Pignoni, M. E. Zuber, X. B. Deng, and Q. X. Zhou for thoughtful discussion and comments. This work was supported by the National Institute of Neurological Disorders and Stroke of the National Institutes of Health under the Award Number R01NS085232 (S.Z.).

## Contribution Statement

R.C., X.L., W.L., and S.Z. conceived the idea, designed the project and approaches. R.C., X.L., W.L., Y.H., carried out the experiments, collected and analyzed the data. R.C. and S.Z. wrote the initial draft of the manuscript. R.C., X.L., W.L., Y.H., and S.Z. revised the manuscript.

## Supplemental information

**Supplemental Table S1.** A list of UAS-RNAi lines used for knocking down mediator complex subunits. All lines were from BDSC except two lines indicated by v#, which were from VDRC.

| Gene | BDSC# | Gene | BDSC# | Gene | BDSC# | Gene | BDSC# |
| --- | --- | --- | --- | --- | --- | --- | --- |
| cdk8 | 35324 | MED13 (Skd) | 34630 | MED11 | 34083 | MED25 | 33595 |
| cdk8 | 67010 | MED1 | 34662 | MED14 | 34575 | MED25 | 42501 |
| cycC | 33753 | MED4 | 34697 | NED15 | 32517 | MED26 | 36829 |
| MED29 | 57259 | MED6 | 33743 | MED16 | 34012 | MED26 | 28572 |
| MED29 | 37545 | MED7 | 34663 | MED17 | 34664 | MED27 | 34576 |
| Kto(MED12) | 34588 | MED7 | 35626 | MED18 | 42634 | MED28 | 32459 |
|  | V#23142 |  |  |  |  |  |  |
|  | V#23143 |  |  |  |  |  |  |
| MED21 | 34731 | MED8 | 34926 | MED20 | 34577 | MED30 | 36711 |
| MED22 | 34573 | MED9 | 33678 | MED23 | 34658 | MED31 | 34574 |
| MED19 | 27559 | MED10 | 34031 | MED23 | 52871 | MED31 | 55220 |
| MED19 | 33710 | MED11 | 57815 | MED24 | 33755 |  |  |

**Figure S1.**
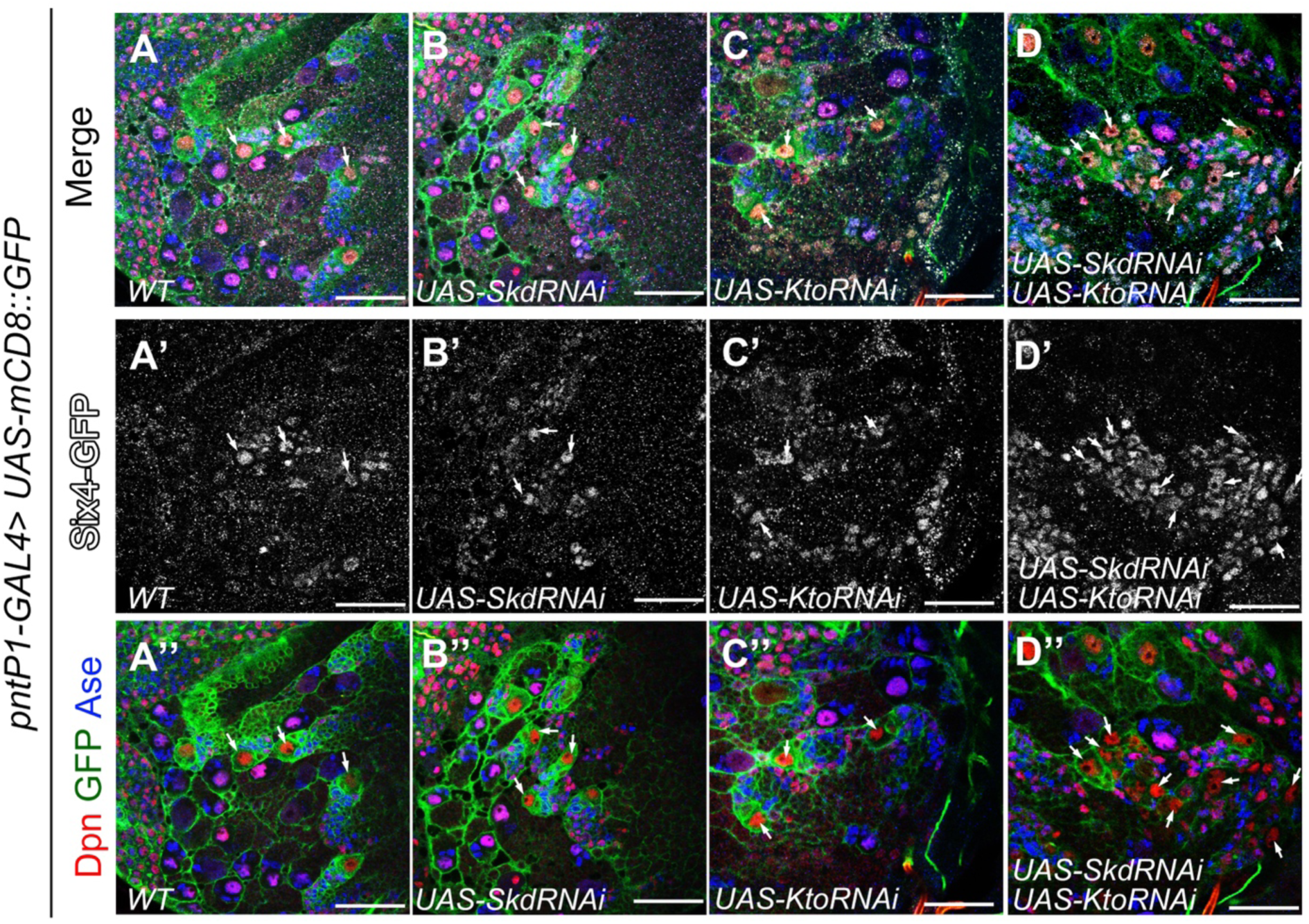
Knockdown of Skd or/and Kto does not affect the expression of Six4 in type II NB lineages. Type II NB lineages are labeled with mCD8-GFP driven by *pntP1-GAL4* and counterstained with anti-Dpn and anti-Ase antibodies. Arrows point to type II NBs. Compared to the expression of Six4-GFP in the wild type brain (A-A”), knockdown of Skd (B-B’), Kto (C-C”), or Skd and Kto together (D-D”) did not obviously affect the expression of Six-GF4 in type II NB lineages.

**Figure S2.**
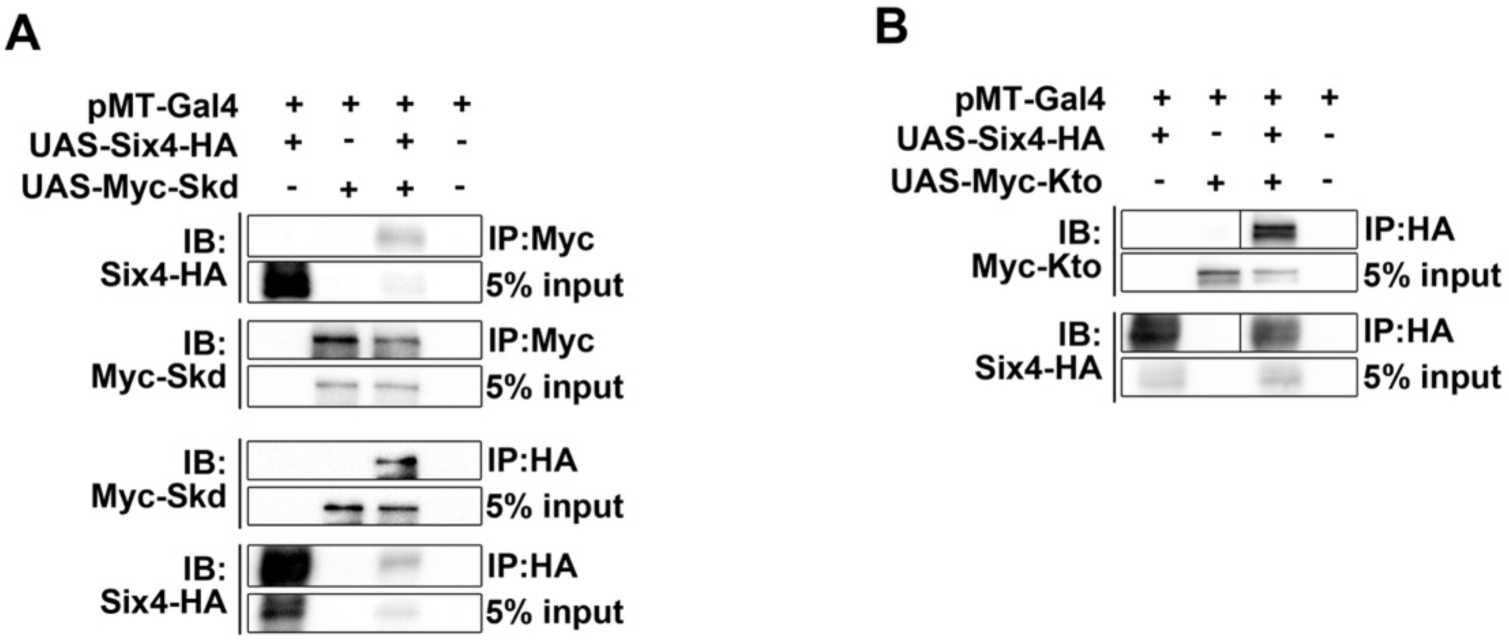
Six4 forms a complex with Skd or Kto. (A-B) Biochemical interactions between Six4-HA and Myc-Skd (A) or between Six4-HA and Myc-Kto (B) are detected by transfecting S2 cells with their corresponding expression constructs followed by co-IP and western blotting analyses.

**Figure S3.**
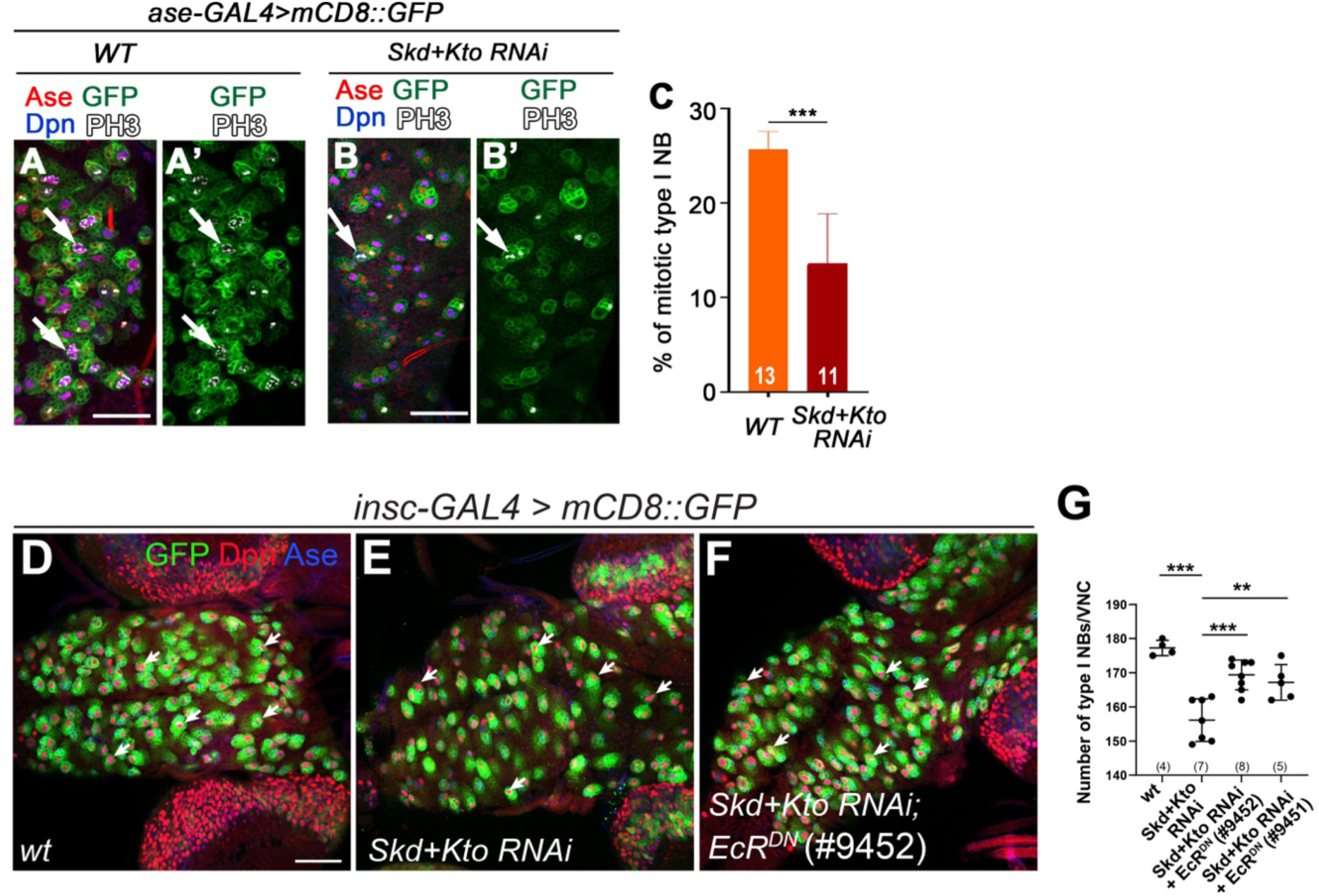
Skd and Kto maintain type I NB self-renewal by antagonizing EcR. In all images, type I NB lineages are labeled with mCD8-GFP driven by *ase-GAL4* (A-B’) or *insc-GAL4* (D-F) and counterstained with the anti-Ase, Dpn, and/or pH3 antibodies. Scale bars: 50µm. (A-B’) pH3 staining in wild type (A-A’) and Skd Kto double knockdown (B-B’) VNCs. Arrows point to mitotic type I NBs. (C) Quantifications of the percentage of pH3+ type I NB in wild type and Skd Kto double knockdown VNCs. Data represent Mean ± SD. ***, *p*<0.001, student *t*-test. The number on each bar represents the number of VNCs examined. (D-F) A wild type VNC (D), and Skd Kto double knockdown VNCs without (E) or with EcR^DN^ expression (F) in type I NBs at 3^rd^ instar larval stage. (G) Quantifications of the number of type I NBs in VNCs with indicated genotypes. Data represent Mean ± SD. ***, *p*<0.001, student *t*-test. The number on each bar represents the number of VNCs examined.

**Figure S4.**
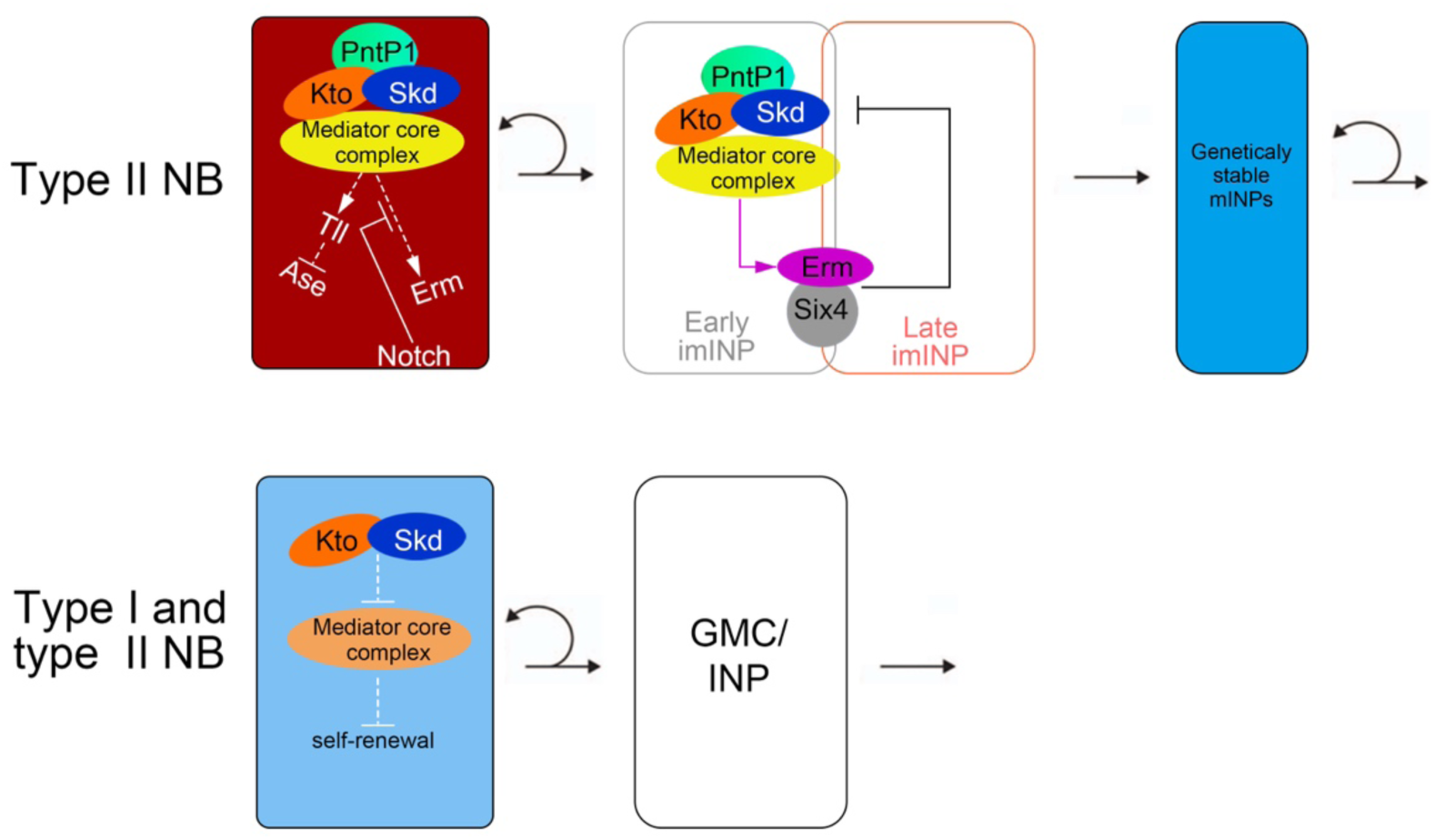
A proposed working model of the functions of Skd/Kto in type II NB lineage development and NB self-renewal. (A) In type II NB lineages, Skd and Kto mediate the activation of different target genes of PntP1 in different cell types. In the NB, Skd and Kto function together with PntP1 to activate Tll, which in turn represses Ase, whereas Notch signaling prevents the activation of Erm by PntP1, Skd and Kto. In early imINPs, Skd and Kto mediate the activation of Erm by PntP1. Erm together with Six4 then promote INP cell fate commitment by inhibiting the activity and expression of PntP1 in late imINPs. Skd and Kto mediate the activation of PntP1 target genes likely by functioning together with a specific subset of subunits of the mediator core complex. (B) In addition to regulating type II NB specification and INP fate commitment, in both type I and type II NBs, Skd and Kto also maintain the self-renewal of NBs by antagonizing the activity of another subset of the mediator core complex subunits that inhibits NB self-renewal.

